# Endogenous ZAP is associated with altered Zika virus infection phenotype

**DOI:** 10.1101/2024.05.23.595518

**Authors:** Nguyen Phuong Khanh Le, Prince Pal Singh, Ahmad Jawad Sabir, Ivan Trus, Uladzimir Karniychuk

## Abstract

The zinc finger antiviral protein 1 (ZAP) has broad antiviral activity. ZAP is an interferon (IFN)-stimulated gene, which itself may enhance type I IFN antiviral response. In a previous study, Zika virus was identified as ZAP-resistant and not sensitive to ZAP antiviral activity. Here, we found that ZAP was associated with the inhibition of Zika virus in Vero cells, in the absence of a robust type I IFN system because Vero cells are deficient for IFN-alpha and -beta. Also, quantitative RNA-seq data indicated that endogenous ZAP is associated with altered global gene expression both in the steady state and during Zika virus infection. Further studies are warranted to elucidate this IFN-alpha and -beta independent anti-Zika virus activity and involvement of ZAP.

## INTRODUCTION

The zinc finger CCCH-type antiviral protein 1 (ZAP) is a cellular protein with broad antiviral activity (1, 2, 3, 4, 5). ZAP targets CpG-rich regions of RNA and ZAP binding to viral RNA mediates its degradation and translational inhibition (3, 6). While ZAP lacks intrinsic RNase activity, it recruits the 5′ and 3′ RNA degradation machinery (7, 8). Using a murine leukemia reporter virus, DDX17 and DHX30 cellular RNA helicases were identified as ZAP co-factors (9, 10). The E3 ubiquitin ligase TRIM25 enhances ZAP activity against alphaviruses (11, 12). ZAP also interacts with the cytoplasmic protein KHNYN to inhibit CpG-enriched human immunodeficiency virus 1 (HIV-1) (13). Recently, Riplet, a protein known for activating the retinoic acid-inducible gene I, was identified as a ZAP co-factor (14). Initially characterized to inhibit murine leukemia virus (15), ZAP has shown efficacy against many viruses, including alphaviruses, filoviruses, influenza virus, porcine reproductive and respiratory syndrome virus, hepatitis B virus, human cytomegalovirus, human T-cell lymphotropic virus type 1, and HIV-1 (1, 6, 16, 17, 18, 19, 20, 21, 22, 23, 24). However, ZAP’s antiviral efficiency is limited against certain RNA viruses, including vesicular stomatitis virus, poliovirus, enterovirus A71, herpes simplex virus type 1, and some flaviviruses (8, 16, 25). Emerging flaviviruses constantly threaten public health. A previous study showed the sensitivity of Japanese encephalitis virus (JEV) to overexpressed ectopic and endogenous ZAP (8). In contrast, yellow fever virus (YFV), dengue virus (DENV), and Zika virus were categorized as ZAP-resistant (8, 16). These are interesting findings because all these flaviviruses have underrepresented CpG content (**Fig S1**) which may help to avoid ZAP recognition (3, 6); however, JEV has the least underrepresented CpG content (**Fig S1**) which supports higher sensitivity to ZAP (8). Here, we further extended studies on ZAP-flavivirus interactions and compared Zika virus infection and transcriptional responses in wild-type and ZAP knockout Vero cells.

## MATERIALS AND METHODS

### Cells

We maintained wild-type Vero E6 cells (ATCC; Vero-ZAP-WT) and ZAP knockout (ZAP-KO) derivatives (Vero-ZAP-KO) in DMEM (Fisher; # 11-965-118) supplemented with 10% heat-inactivated fetal bovine serum (FBS, Fisher; #A5256801), 1x Penicillin-Streptomycin (Fisher; #15140122), and 2.67 mM sodium bicarbonate (Fisher; # 25080094) at +37°C in a 5% CO_2_ humidified incubator. To generate ZAP-KO cell line, we used the guide RNA (gRNA) GTCTCTGGCAGTACTTGCGA which targets the first exon of *Chlorocebus sabaeus* (Gene ID: 103226990) ZC3HAV1 gene, which is required for the expression of all ZAP isoforms. We also used a non-targeting control gRNA ACGGAGGCTAAGCGTCGCAA for control cells. These gRNAs were transiently transfected into cells using GenCrispr NLS-Cas9-NLS Nuclease (GenScript). Subsequently, transfected cells were seeded in 96-well plates through limiting dilution to generate isogenic single clones. Following expansion, the clones were genotyped via Sanger sequencing, employing primers (ATCGCTGGGCTGGACTAACG, GCAGAGAAGGGAGTGGCTGAA) to identify indels (location of indels on *Chlorocebus sabaeus* genome: −2bp deletion, NW_023666072.1: 4677550-4677551). The knockout subclone verified by genotyping was further confirmed by western blot for ZAP expression (**Fig 1A, Fig S2**) as described below. The negative control cells were validated by Sanger sequencing to confirm there were no indels. We confirmed the absence of mycoplasma contamination in cells using the LookOut Mycoplasma PCR Detection Kit (Millipore Sigma).

**Fig 1.**
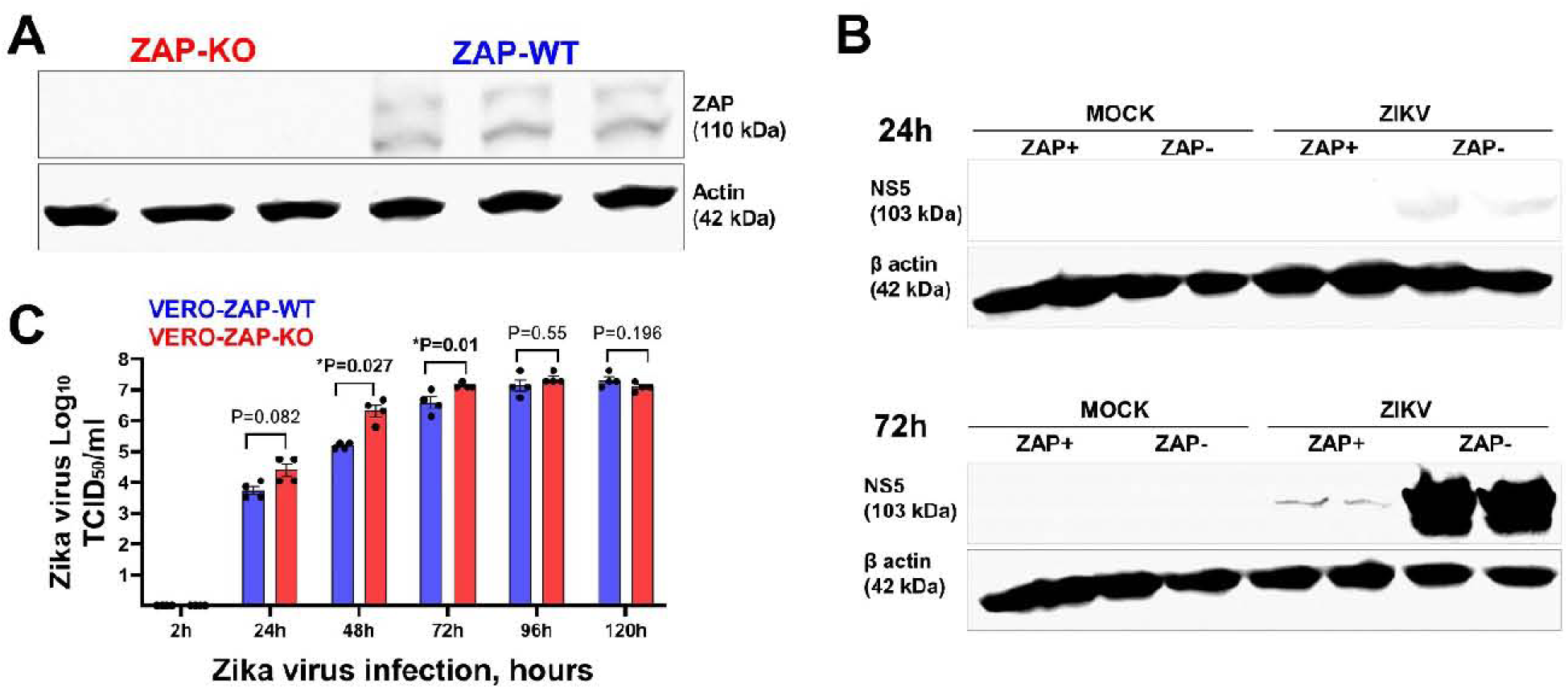
Infection phenotypes of Zika virus in wild-type (Vero-ZAP-WT) and ZAP knockout (Vero-ZAP-KO) cells. (**A**) ZAP Western blot. KO status was also confirmed by Sanger sequencing. Rabbit anti-ZAP IgG Abs (1:1,000 dilution; ANTI-ZC3HAV1; #HPA059096-100UL, Sigma-Aldrich), with the specific ZAP band appearing at 110 kDa, were used. Beta-actin was used as a western blot loading control (42 kDa). A normalized 50 µg of protein was used for all samples. Three replicates. Multiple bands may represent isoforms described for human ZAP (3) and still not experimentally characterized for monkeys. The full blot image is in **Figure S2**. (**B**) Zika virus NS5 protein Western blot. Reduced Zika virus NS5 protein loads in Vero-ZAP-WT cells as compared to Vero-ZAP-KO cells at 24 h and 72 h after inoculation, MOI 1. Rabbit anti-Zika virus NS5 IgG Abs (1:1,000 dilution; GTX133312, GeneTex), with the specific NS5 band appearing at 103 kDa, were used. Beta-actin was used as a western blot loading control (42 kDa). A normalized 50 µg of protein was used for all samples. Two biological replicates for each condition are shown. The full blot image is in **Figure S3**. (**C**) Zika virus infectious titers in supernatants of Vero-ZAP-WT and Vero-ZAP-KO cells at different times after inoculation, MOI 1. The experiment was done in four replicates for each cell line and sampling time. Supernatants from all mock-infected samples were negative. Statistics—an unpaired t-test. *A P-value of < 0.05 was considered statistically significant.

### Viruses

We utilized the Asian lineage Zika virus H/PF/2013 strain [GenBank: KJ776791.2]. Zika virus was rescued using reverse genetics (26). Subsequently, the virus was propagated in Vero cells cultured in DMEM supplemented with 2% FBS and 1% penicillin/streptomycin. Supernatants were harvested, centrifuged (12,000 g, 20 min, +4°C), aliquoted, and frozen (−80°C). Infectious virus titers were quantified using Vero cells via and endpoint dilution assay. The fifty percent tissue culture infective dose (TCID_50_) endpoint titers were calculated as previously described (26, 27, 28, 29). Media from virus-negative Vero cells were processed in the same manner as the virus stocks and used as control. The absence of mycoplasma contamination in the virus stocks was confirmed using the LookOut Mycoplasma PCR Detection Kit (Millipore Sigma; #MP0035).

### Zika virus infection kinetics

Vero-ZAP-WT and Vero-ZAP-KO cells were seeded in 6-well plates in DMEM supplemented with 10% inactivated fetal bovine serum for 24 h. Four wells for each cell line, representing four biological replicates (cells from different passages and flasks in each of 4 wells), were inoculated with Zika virus at a multiplicity of infection (MOI) of 1. After 2 h at 37°C, the inoculum was removed, the cells were washed three times, covered with 3 ml of DMEM supplemented with 2% FBS, and maintained at 37°C. At 2, 24, 48, 72, 96, and 120 h post-inoculation, 150 µl of media supernatants were collected, cleared by centrifugation (2,000 g, 10 min), and frozen at - 80°C for the end-point dilution TCID_50_ assay (26, 29, 30, 31, 32, 33).

### Western blot

Untreated, mock-infected, or Zika-infected (MOI 1) Vero-ZAP-WT and Vero-ZAP-KO cells were washed, pelletized, and lysed in 100 μL of RIPA buffer (Fisher; # PI89900) supplemented with 1× Halt protease inhibitor cocktail (Fisher; #PI87786) and 1× EDTA (Fisher; #BP2473100). The lysates were then incubated on wet ice for 10 minutes, gently vortexed, and centrifuged (12,000g, 10 min, +4 °C). Supernatants were aliquoted into prechilled low protein-binding (Sarstedt; #72.706.600) and stored at −80°C. After quantifying protein concentration with the Pierce BCA Protein Assay Kit (Fisher; # PI23227), 50 μg of protein extract was diluted in 4× reducing Laemmli SDS sample buffer (Fisher; #AAJ60015AC), heated at 95°C for 10 min, and resolved on 10% Mini-PROTEAN TGX precast protein gels (Bio-Rad; # 4561035) in 1× Tris/Glycine/SDS running buffer (Bio-Rad; #1610732) at a voltage of 125V for 70 min. Proteins were transferred on methanol-activated 0.45 μm Immun-Blot Low Fluorescence polyvinylidene fluoride (PVDF) membranes using the Trans-Blot Turbo RTA Mini 0.45 µm LF PVDF Transfer Kit (Bio-Rad; #1704274). The PVDF membranes were blocked with 1× VWR Life Science Rapidblock Blocking Solution (VWR; #97064-124) for 1 h at room temperature. Subsequently, the membranes were incubated overnight at +4°C with primary antibody diluted in blocking solution. The membranes were washed four times with TBST (1× Bio-Rad Tris Buffered Saline containing 0.1% Tween-20) and then incubated with secondary antibody, along with hFAB Rhodamine Anti-Actin Primary Antibody (Recombinant human β-actin), diluted in Blocking Solution for 1 h at room temperature, and washed four times with TBST. Fluorescent blots were imaged on the ChemiDoc MP Imaging system using an appropriate detection channel with Image Lab Touch Software (Bio-Rad).

We used antibodies (Abs): Rabbit anti-ZAP IgG Abs (1:1,000 dilution; ANTI-ZC3HAV1; #HPA059096-100UL, Sigma-Aldrich), Rabbit anti-Zika virus NS5 IgG Abs (1:1,000 dilution; #GTX133312, GeneTex), hFAB Rhodamine Anti-Actin Primary Abs (1:2,500; #12004164, Bio-Rad), and IRDye 680RD Goat anti-Rabbit IgG secondary Abs (1:10,000; #926-68071; LI-COR).

### Comparative RNA-seq analysis in wild-type and ZAP-KO cells

Vero-ZAP-WT and Vero-ZAP-KO cells were seeded at 6×10^5^ cells per well in 6-well plates, with 4 plates per cell line, in DMEM supplemented with 10% FBS for 24 h. Wells in two plates from each cell line were either mock-inoculated with media from virus-negative cells (4-6 well replicates for Vero-ZAP-WT and Vero-ZAP-KO) or inoculated with MOI 1 of Zika virus (6 well replicates for Vero-ZAP-WT and Vero-ZAP-KO). After 2 h at 37°C, the inoculum was removed, cells were washed, and covered with DMEM supplemented with 2% FBS, then maintained at 37°C. At 24 h and 72 h post-inoculation, media was removed, cells were washed with commercial 1× PBS, homogenized in 1 mL of TRI Reagent Solution (Fisher; # AM9738), and RNA was extracted according to the manufacturer’s protocol using the PureLink RNA Mini Kit (Fisher; 12-183-018A). A total of 4-6 replicate samples were obtained for mock-inoculated Vero-ZAP-WT, mock-inoculated Vero-ZAP-KO, Zika-infected Vero-ZAP-WT, and Zika-infected Vero-ZAP-KO cells, at both 24 and 72 h. Zika infection was confirmed by virus-specific RT-qPCR with a previously validated SYBR one-step RT-qPCR assay (26, 28, 34). RNA quantification was performed using the Qubit RNA High Sensitivity kit (Fisher; # Q32855) and Qubit 2.0 Fluorometer (Fisher). RNA quality was assessed using a bioanalyzer (Agilent Technologies 2100) to ensure all samples had RNA Integrity Number (RIN) values above 9.0. DNA contamination was removed using the TURBO DNA-free Kit (Fisher; # AM1907). mRNA with intact poly(A) tails was enriched using the NEBNext Poly(A) mRNA Magnetic Isolation Module (NEB; # E7490L) and utilized for library constructions with the NEBNext Ultra II Directional RNA Library Prep Kit for Illumina along with NEBNext Multiplex Oligos for Illumina (96 Unique Dual Index Primer Pairs; NEB; # E7760L). The libraries were sequenced on the NovaSeq platform as paired-end reads using the S1 v1.5 kit with 300 cycles (Illumina). Raw sequencing data were deposited to NCBI BioProject (PRJNA1101086).

FASTQ files were trimmed for adaptor sequences and filtered for low-quality reads using *Trimmomatic*. RNA-seq analysis was conducted as previously described (29, 30, 33, 35) with modifications. Briefly, paired-end reads were processed using the *Kallisto* pseudo-alignment method (36) and quantified by mapping to a transcripts database generated from the African Green Monkey (vervet) reference genome assembly (ENSEMBL *Chlorocebus sabaeus* 1.1) (37). The count table for RNA-seq data was assembled using the *tximport::tximport* function in R. After importing data to the R environment we removed data for genes with low expression using the *edgeR::filterByExpr* function (38). Normalization was then performed using the *edgeR::calcNormFactors* function, and the *limma::voom* function was used to convert the data into a normal distribution. We calculated differential expression using the *limma::lmFit* function with empirical Bayes moderation via the *limma::eBayes* function (**Table S1**).

Several experimental conditions were applied to compare Differentially Expressed Genes (DEGs) (**Table S1**). First, we compared gene expression directly between mock-infected ZAP-WT and ZAP-KO Vero cells. We then used two strategies to examine transcriptional responses during Zika virus infection in wild-type and knockout cells: The first strategy involved a direct comparison of gene expression between Zika virus-infected ZAP-WT and ZAP-KO Vero cells. The second strategy compared transcriptional responses between non-infected and infected Vero-ZAP-WT cells, followed by a similar comparison between non-infected and infected Vero-ZAP-KO cells. Finally, significantly affected genes (FDR < 0.05 and log2 FC > 1) were summarized in **Table S2.**

Significantly affected DEGs from each experimental condition were further analyzed by comparing these DEGs (**Table S2**) to a set of 194 genes involved in the interferon signaling pathway. This Interferon Signaling Gene Set was obtained from Harmonizome 3.0 (https://maayanlab.cloud/Harmonizome/gene_set/Interferon+Signaling/Reactome+Pathways) and is derived from the Reactome Pathways dataset (https://reactome.org/PathwayBrowser/#/R-HSA-913531). The DEGs significantly affected in each condition that are part of the interferon signaling pathway were summarized in **Table S3** and visualized in **Fig 2** for pairwise comparison.

**Fig 2.**
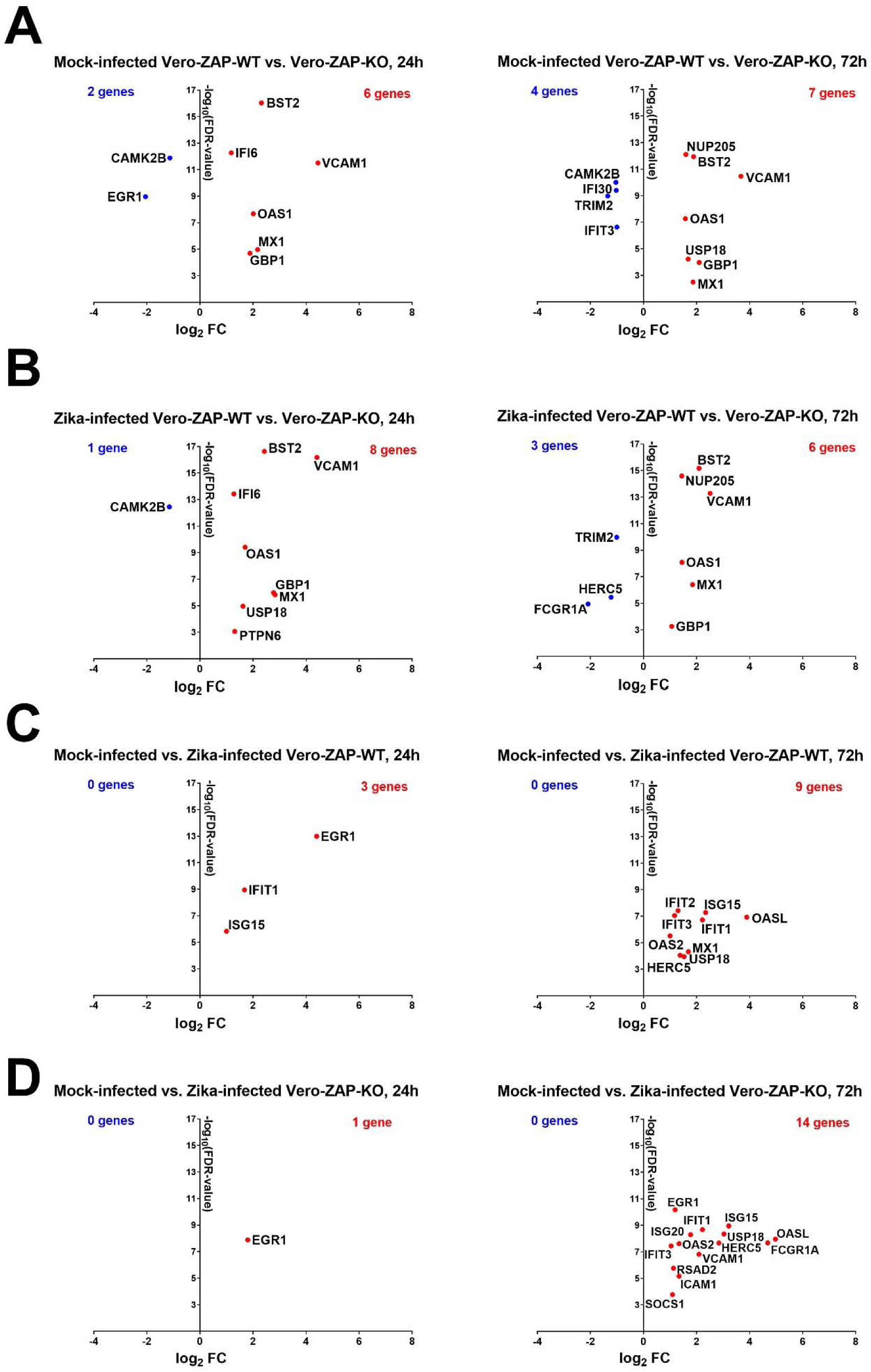
Significantly affected interferon signaling genes in different experimental conditions. (**A**) Mock-infected Vero-ZAP-WT versus Vero-ZAP-KO cells. Direction of analysis: WT/KO. **Tables S3B, C**. (**B**) Zika-infected Vero-ZAP-WT versus Vero-ZAP-KO cells. Direction of analysis: WT/KO. **Tables S3D, E.** (**C**) Mock-infected versus Zika-infected Vero-ZAP-WT cells. Direction of analysis: Zika-infected/Mock-infected. **Tables S3F, G.** (**D**) Mock-infected versus Zika-infected Vero-ZAP-KO cells. Direction of analysis: Zika-infected/Mock-infected. **Tables S3H, I.** Plots of the upregulated (red) and downregulated (blue) genes. **FDR**: The false discovery rate. Genes with FDR < 0.05 and log_2_ fold change (FC) > 1 were significantly affected.

In addition to pairwise comparison, we performed a gene set enrichment analysis using the *limma::camera* function for each experimental condition. The gene set enrichment analysis has higher sensitivity than pairwise comparison because not only genes with FDR < 0.05 and log_2_ fold change (FC) > 1 (**Table S2**), but each gene in the entire input set (**Table S1)** is statistically processed and used for annotation (39, 40). Gene set enrichment analysis was conducted using Gene Ontology (GO) annotation for *Chlorocebus sabaeus* obtained from g:Profiler (41), and significant (FDR < 0.25) GO processes for each experimental condition were summarized in **Table S2**.

### Statistics

Zika infectious TCID_50_ titers in Vero-ZAP-WT and Vero-ZAP-KO cells were compared using an unpaired t-test; log_10_ transformation was performed on the TCID_50_ values before the t-test. A *P*-value < 0.05 was considered statistically significant.

## RESULTS

Zinc finger antiviral protein (**Fig 1A; Fig S2**) affected expression of Zika virus NS5 protein, as determined by Western blot at 24 h and 72 h (**Fig 1B; Fig S3**). Specifically, at 24 h post-inoculation, faint Zika NS5 bands were visible in Vero-ZAP-KO cells, while the assay sensitivity was insufficient to detect NS5 in Vero-ZAP-WT cells. By 72 h post-inoculation, Zika NS5 was detected in both Vero-ZAP-KO and Vero-ZAP-WT cells, with protein loads considerably higher in Vero-ZAP-KO cells (**Fig 1B**). In addition to NS5 Western blot, infectious Zika virus titers in the supernatants of Vero-ZAP-KO cells were significantly higher than Vero-ZAP-WT titers at 48 and 72 hours (**Fig 1C**).

For RNA-seq analysis, we first compared global gene expression between mock-infected Vero-ZAP-WT and Vero-ZAP-KO cells. A total of 758 and 992 genes were significantly affected and differentially expressed between the two cell lines at 24 and 72 h, respectively (**Tables S2A, B**). Second, we compared gene expression between Zika-infected Vero-ZAP-WT and Vero-ZAP-KO cells. At 24 h, we identified 631 genes that were significantly affected and differentially expressed (**Table S2E**). This number increased to 864 genes at 72 h (**Table S2F**). When we compared significant DEGs between steady state (mock-infected) (**Tables S2A, B**) and Zika virus-infected conditions (**Tables S2E, F**), 89 (14.1%) and 196 (22.7%) genes were uniquely affected during infection at 24 h and 72 h (**Tables S2E, F**). Thus, endogenous ZAP influences global gene expression both under steady-state conditions and during Zika virus infection.

Next, we compared global gene expression between mock-infected and Zika-infected Vero-ZAP-WT cells. A total of 40 and 117 genes were significantly affected by infection at 24 h and 72 h (**Tables S2I, J**). Finally, we compared global gene expression between mock-infected and Zika-infected Vero-ZAP-KO cells. A total of 14 and 342 genes were significantly affected by infection at 24 h and 72 h (**Tables S2M, N**). The higher number of significantly affected genes in infected ZAP-KO cells (342 genes) compared to ZAP-WT cells (117 genes) at 72 h is not surprising. This is probably because ZAP, in addition to its antiviral activity, is known to bind cellular mRNA and affect gene expression leading to the resolution of antiviral and immune transcriptional responses (42, 43).

Next, we investigated whether ZAP, beyond its impact on global gene expression, specifically affects genes involved in the interferon signaling pathway. To do this, we compared significantly affected DEGs in different experimental conditions (**Table S2**) with a set of 194 genes involved in the interferon signaling pathway (**Table S3A**). In mock-infected Vero-ZAP-WT cells compared to mock-infected Vero-ZAP-KO cells, 8 DEGs were significantly affected at 24 h and 11 DEGs at 72 h (**Fig 2A; Table S3B, C**). Most affected DEGs were overexpressed in Vero-ZAP-WT. Interestingly, during Zika virus infection (**Fig 2B; Table S3D, E**), endogenous ZAP had minimal impact on the expression of interferon signaling pathway genes because the pattern of significantly affected genes in Zika virus-infected ZAP-WT and ZAP-KO cells was very similar to that seen in mock-infected cells (**Fig 2A, B; Table S3B-E**). However, when comparing Zika virus-infected cells to mock-infected cells, both ZAP-WT and ZAP-KO cells showed increased responses to infection, including overexpression of EGR1, IFIT1, ISG15, ISG20, IFIT2, OASL, ICAM1, OAS2, RSAD2, and SOCS1 (**Fig 2C, D; Table S3F-I**) which were not identified by other analytical strategies (**Fig 2A, B; Table S3B-E**).

To further extend analysis beyond pairwise comparisons of top DEGs (**Fig 2; Table S3**), we conducted GO analysis. This method offers high sensitivity due to its robust annotation framework, standardized terminology, hierarchical structure, integration with biological knowledge, continuous updates, and statistical rigor where not only genes with FDR < 0.05 and log_2_ fold change (FC) > 1 (**Table S2**), but each gene in the entire input set (**Table S1)** is statistically processed and used for annotation (39, 40).

Mock-infected Vero-ZAP-WT cells when compared to mock-infected Vero-ZAP-KO cells had 92 (24 h) and 178 (72 h) significantly affected GO biological processes (**Tables S2C, D**). However, when we searched among affected GO biological processes keywords—“immun*” and "viral/virus,” we did not find affected processes containing these keywords (**Tables S2C, D**). This contrasts with affected GO biological processes in infected cells described below, suggesting the specificity of findings in Zika-infected cells.

Zika-infected Vero-ZAP-KO cells had 192 (24 h) and 481 (72 h) significantly affected GO biological processes when compared to Zika-infected Vero-ZAP-WT cells (**Tables S2G, H**). Zika-infected Vero-ZAP-WT cells had total 303 (24 h) and 864 (72 h) affected GO biological processes as compared to mock-infected ZAP-WT cells (**Tables S2K, L**). Zika-infected Vero-ZAP-KO cells had total 287 (24 h) and 384 (72 h) affected GO biological processes as compared to mock-infected ZAP-WT cells (**Tables S2O, P**).

We narrowed down analysis and compared GO biological processes related to immune responses in mock-infected Vero-ZAP-WT and Vero-ZAP-KO cells at 24 and 72 h. As expected, no affected “immune*” GO processes were detected in mock-infected control cells (**Table S2C, D**). After direct comparison between Zika-infected Vero-ZAP-KO and Vero-ZAP-WT cells, we found one significantly affected immune process at 24 h and 72 h (**Fig 3A; Tables S2G, H**). At each time point immune GO processes were upregulated in ZAP-WT cells. When compared to mock-infected Vero-ZAP-WT cells, infected Vero-ZAP-WT cells showed one significantly affected process at 24 h, and the number increased at 72 h reaching 16 affected processes (**Fig 3B; Tables S2K, L**). When compared to mock-infected Vero-ZAP-KO cells, infected Vero-ZAP-KO cells showed one significantly affected process at 24 h, and the number increased at 72 h reaching only 4 affected processes (**Fig 3C; Tables S2O, P**).

**Fig 3.**
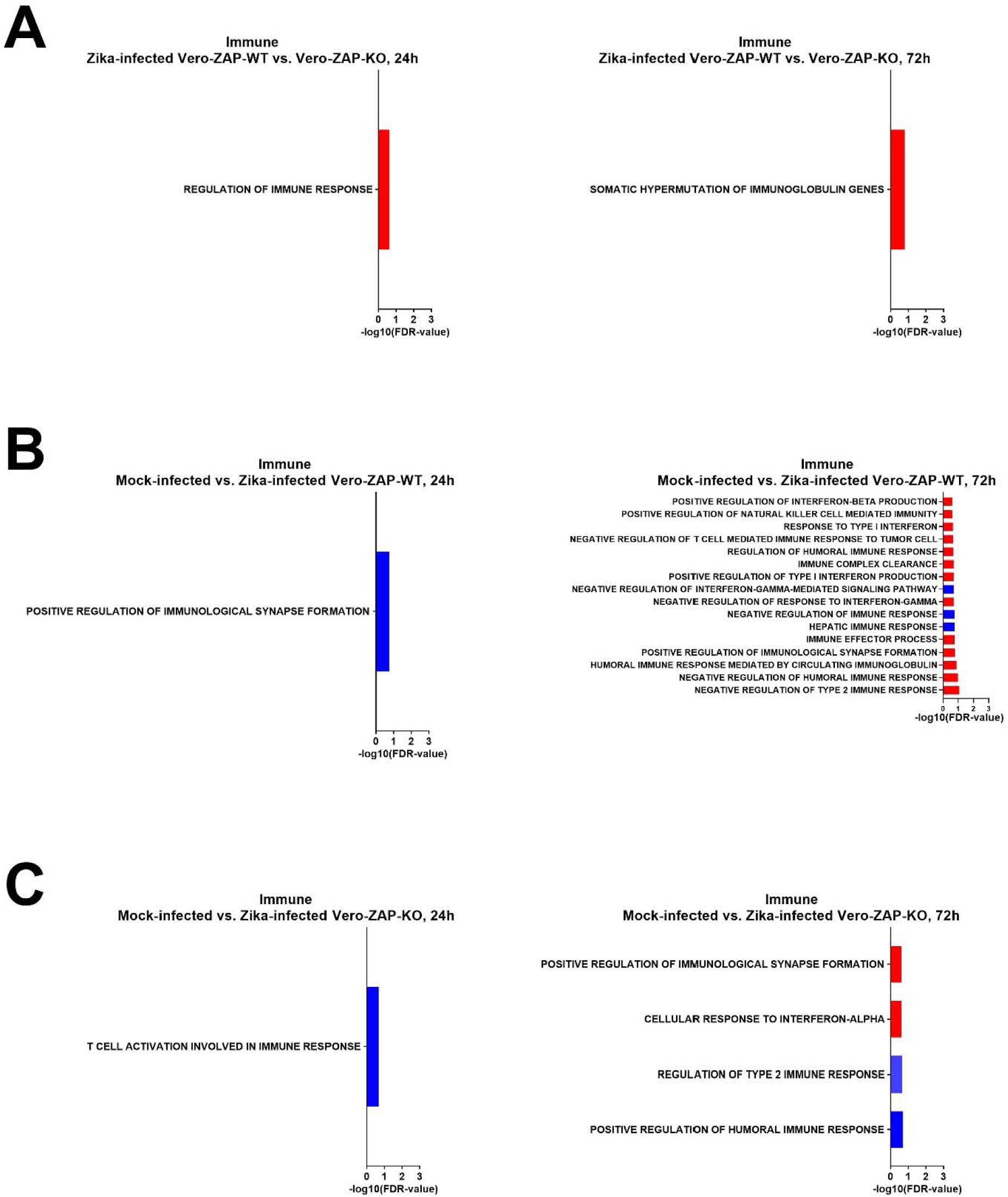
Biological “immune*” processes significantly affected in Vero-ZAP-WT and Vero-ZAP-KO cells at 24 h and 72 h after Zika virus infection. (**A**) Zika-infected Vero-ZAP-WT versus Vero-ZAP-KO cells. Direction of analysis: WT/KO. **Tables S2G, H.** (**B**) Mock-infected versus Zika-infected Vero-ZAP-WT cells. Direction of analysis: Zika-infected/Mock-infected. **Tables S2K, L.** (**C**) Mock-infected versus Zika-infected Vero-ZAP-KO cells. Direction of analysis: Zika-infected/Mock-infected. **Tables S2O, P.** Red bars–upregulated processes. Blue bars– downregulated processes. **FDR**: The false discovery rate.

The similar trend was observed when we narrowed down analysis and compared GO biological processes related to virus infection with “viral/virus” keywords (**Tables S2C, D, G, H, K, L, O, P**).

## DISCUSSION

In a previous study, Zika virus was identified as ZAP-resistant and not sensitive to ZAP antiviral activity (8). Here, we showed that ZAP acts as restriction factor for Zika virus in Vero cells. Also, quantitative RNA-seq data indicated that endogenous ZAP is associated with altered global gene expression both in the steady state and during Zika virus infection.

The previous study in A549 cells showed JEV sensitivity to ectopic and endogenous ZAP (8). In the same study, Zika virus was not sensitive to ectopic ZAP; the sensitivity of Zika virus to endogenous ZAP was not tested (8). Another study using ZAP wild-type and ZAP knockout A549 cells showed that wild-type Zika virus was also not sensitive to antiviral ZAP effects (44). Here, however, we showed that Zika virus was sensitive to endogenous ZAP in Vero cells. Specifically, we observed reduced infectious titers and Zika virus NS5 protein expression in wild-type cells compared to ZAP knockout cells. This discrepancy may be due to the different cell lines used: previous studies utilized A549 cells derived from human lung cancer tissue (8), while we used Vero cells from a healthy monkey. Antiviral signaling for many cellular proteins can be highly cell-specific (45). Another factor is the species difference: the previous study used a human cell line, and we used an African green monkey cell line, a possible natural reservoir host of Zika virus (46). Species-specific ZAP effects are documented (47, 48), but comparative studies on ZAP antiviral activity between primates are lacking.

Using pairwise comparisons of DEGs, we found that endogenous ZAP impacts global gene expression both in the steady state and during Zika virus infection. ZAP, however, had minimal impact on the genes expression of the interferon signaling pathway. Interestingly, Zika virus infection in both ZAP-WT and ZAP-KO cells was associated with increased expression of genes involved in the interferon signaling pathway (**Figs 2C, D**). These findings are interesting and novel in the context of Zika infection because Vero cells are deficient in IFN-α/β (4, 49). We do not know yet how IFN-α/β-independent anti-Zika transcriptional responses work in Vero cells; however, these responses may be attributed to type III IFN lambda. Indeed, IFN lambda 1 was overexpressed in both infected ZAP-WT and ZAP-KO cells as compared to corresponding mock-infected control cells at 72 h (**Table 1; Tables S2J, N**). Comparison of Zika virus-infected ZAP-WT versus ZAP-KO cells suggests that ZAP may downregulate IFN lambda 1 expression (FDR = 0.0005 and log_2_ FC = −1.006; **Table S2F**), which is supported by ability of ZAP to bind cellular mRNAs and affect gene expression leading to the resolution of antiviral and immune transcriptional responses (42, 43).

**Table 1.** Interferon lambda gene expression during Zika infection in Vero-ZAP-WT and Vero-ZAP-KO cells.

|  |  | Gene symbol | Gene name | Log<sub>2</sub> FC | FDR |
| --- | --- | --- | --- | --- | --- |
| ZAP-WT* | 72 h | IFNL1 | interferon lambda 1 | 4.82 | 3.83E-08 |
| ZAP-KO* | 72 h | IFNL1 | interferon lambda 1 | 5.13 | 1.52E-10 |

**FC:** Fold change. **FDR:** False discovery rate. Data are in **Tables S2J, N**. \*IFNL1 expression during Zika infection in ZAP-WT and ZAP-KO Vero cells compared to mock-infected ZAP-WT and ZAP-KO cells, respectively. Direction of analysis: Zika-infected/Mock-infected.

Interferon lambda induces antiviral response activating the same intracellular signaling pathways and have similar biological activities as IFN-α/β, but signal via distinct receptor complex (50). Also, it is known that Vero cells respond to hantavirus infection by secreting abundantly IFN lambda (51), and IFN lambda protects the female reproductive tract against Zika virus infection (52).

While pairwise comparisons of DEGs showed minimal impact of ZAP on the gene expression of the interferon signaling pathway, more sensitive GO analysis suggests that ZAP may augment transcriptional antiviral responses. Specifically, when Zika virus-infected ZAP-KO and ZAP-WT cells were compared to their corresponding mock-infected control cells, ZAP-WT infected cells showed 4 times more significantly affected GO “immune” processes than ZAP-KO infected cells at 72 h (16 versus 4 processes, **Fig 3B, C; Tables S2L, P**). ZAP is an IFN-stimulated gene, which itself may enhance type I IFN antiviral response (1, 2, 3, 4, 5, 43, 53, 54). We do not know however how ZAP may enhance type I IFN antiviral response in IFN-α/β-deficient Vero cells, suggesting further research on ZAP and type III IFN interactions.

In this study, we used the Asian Zika virus strain, which is related to strains responsible for the 2015 Zika epidemic. Historical African Zika strains are known to cause more aggressive infections and elicit stronger and faster innate immune responses (28, 31, 55). Previous studies on Zika-ZAP interactions also used the Asian strain (8, 44). Therefore, it would be interesting to conduct comparative studies to investigate how ZAP influences infections caused by both recent and historical Zika strains across different cell lines.

Our study is confined to a single Vero-ZAP-KO clone, and conducting additional tests using multiple clones could yield further insights. Such an endeavor falls beyond the scope of this brief report; however, our RNA-seq results find indirect support from previous research conducted on different uninfected wild-type and ZAP-KO cell lines. Specifically, global gene expression analysis with Agilent microarrays revealed 1,065 upregulated and 776 downregulated transcripts in uninfected wild-type and ZAP-KO HeLa cells (42). Another study using RNA-seq identified a number of DEGs between uninfected ZAP-WT and ZAP-KO A549 cells (56). In the most recent study, the authors demonstrated that ZAP influences hundreds of transcripts in ZAP-WT and ZAP-KO HEK cells (57). Directly comparing the DEGs between our study and the above studies poses challenges due to variations in cell types, experimental design, and the employed bioinformatics. Nonetheless, the overarching trend in all previous studies strongly corroborates significant transcriptional differences between uninfected ZAP-WT and ZAP-KO Vero cells (**Tables S2A-D**).

The above studies did not use virus infection for RNA-seq comparison between ZAP-WT and ZAP-KO cells. Thus, to our knowledge, our study provides the first high-throughput RNA-seq dataset highlighting the impact of ZAP on global gene expression during virus infection, at least in Vero cells. During submission of this manuscript, a study indirectly supporting our findings was published (57). The authors used RNA-seq in ZAP-WT and ZAP-KO HEK cells transfected with viral mimetic RNA oligos and showed that ZAP has a role in transcriptome regulation in basal and antiviral states, which is in accordance with findings in the present study.

Altogether, we found that ZAP was associated with the inhibition of Zika virus in the absence of a robust type I IFN system (Vero cells are deficient for IFN-α/β). Enrichment analysis of GO processes also suggests the potential importance of ZAP in augmenting transcriptional responses against Zika virus even in the absence of IFN-α/β. Further investigations are warranted to elucidate the mechanisms underlying these IFN-α/β-independent antiviral cellular responses to Zika virus infection, and potential antiviral interactions between ZAP and type III IFNs.

## LIST OF ABBREVIATIONS

ZAP: zinc finger antiviral protein
IFN: interferon
WT: wild-type
KO: knockout
FC: fold change
FDR: false discovery rate
NS5: non-structural protein 5
RT-qPCR: reverse transcription quantitative polymerase chain reaction
TCID_50_: fifty percent tissue culture infective dose
JEV: Japanese encephalitis virus
YFV: yellow fever virus
DENV: dengue virus
HIV-1: human immunodeficiency virus 1
ZIKV: Zika virus
WNV: West Nile virus

## CONSENT FOR PUBLICATION

Not applicable.

## ETHICS APPROVAL AND CONSENT TO PARTICIPATE

Not applicable.

## AVAILABILITY OF DATA AND MATERIALS

Data are provided within the manuscript or supplementary information files. Raw RNA-seq files are deposited to NCBI BioProject (PRJNA1101086).

## AUTHORS’ CONTRIBUTIONS

Conceptualization: UK. Investigation: NPKL, PPS, AJS, UK. Data analysis: NPKL, PPS, IT, UK. Funding: UK. Resources: UK. Writing—original draft preparation: NPKL, UK. Writing—review and editing: UK, NPKL, IT, PPS.

## SUPPLEMENTARY TABLES

**Table S1** Raw RNAseq data (DEGs).

**Table S2** Significantly affected DEGs and GO processes.

**Table S3** Expression of Interferon Signaling Genes in different experimental conditions.

**Fig S1. CpG dinucleotide content in flavivirus genomes.** The bar plot is the CpG frequency in the whole flavivirus RNA genomes (the ratio of observed and arithmetically expected CpG dinucleotides determined by SSE 1.4 software). Line graphs show the number of CpG dinucleotides within the RNA genome with a 50 nt sliding window. Calculations were made with an R script (https://github.com/itrus/Mapping-ROIs-in-genetic-sequences/blob/main/CpG-map-linear.R). **JEV**—Japanese Encephalitis virus (GenBank: MK585066.1 reference genome); **ZIKV**—Zika virus (GenBank: KJ776791.2); **DENV**—dengue virus (GenBank: U87411.1); **YFV**—yellow fever virus (GenBank: X03700.1); **WNV**—West Nile virus (GenBank: DQ211652.1).

**Fig S2. ZAP Western blot.** The full blot image. Rabbit anti-ZAP IgG Abs (1:1,000 dilution; ANTI-ZC3HAV1; #HPA059096-100UL, Sigma-Aldrich), with the specific ZAP band appearing at 110 kDa, were used. Multiple bands may represent four isoforms described for human ZAP (3) and still not experimentally characterized for monkeys.

**Fig S3. Zika virus NS5 Western blot.** The full blot image. Rabbit anti-Zika virus NS5 IgG Abs (1:1,000 dilution; GTX133312, GeneTex), with the specific NS5 band appearing at 103 kDa, were used.

## REFERENCES

1. Nguyen LP, Aldana KS, Yang E, Yao Z, Li MMH. Alphavirus Evasion of Zinc Finger Antiviral Protein (ZAP) Correlates with CpG Suppression in a Specific Viral nsP2 Gene Sequence. Viruses. 2023;15(4).

2. MacDonald MR, Machlin ES, Albin OR, Levy DE. The zinc finger antiviral protein acts synergistically with an interferon-induced factor for maximal activity against alphaviruses. J Virol. 2007;81(24):13509–18.

3. Li MMH, Aguilar EG, Michailidis E, Pabon J, Park P, Wu X, et al. Characterization of Novel Splice Variants of Zinc Finger Antiviral Protein (ZAP). J Virol. 2019;93(18).

4. Crosse KM, Monson EA, Beard MR, Helbig KJ. Interferon-Stimulated Genes as Enhancers of Antiviral Innate Immune Signaling. J Innate Immun. 2018;10(2):85–93.

5. Hayakawa S, Shiratori S, Yamato H, Kameyama T, Kitatsuji C, Kashigi F, et al. ZAPS is a potent stimulator of signaling mediated by the RNA helicase RIG-I during antiviral responses. Nat Immunol. 2011;12(1):37–44.

6. Takata MA, Goncalves-Carneiro D, Zang TM, Soll SJ, York A, Blanco-Melo D, et al. CG dinucleotide suppression enables antiviral defence targeting non-self RNA. Nature. 2017;550(7674):124-7.

7. Zhu Y, Gao G. ZAP-mediated mRNA degradation. RNA Biol. 2008;5(2):65–7.

8. Chiu HP, Chiu H, Yang CF, Lee YL, Chiu FL, Kuo HC, et al. Inhibition of Japanese encephalitis virus infection by the host zinc-finger antiviral protein. PLoS Pathogens: Public Library of Science; 2018.

9. Ye P, Liu S, Zhu Y, Chen G, Gao G. DEXH-Box protein DHX30 is required for optimal function of the zinc-finger antiviral protein. Protein Cell. 2010;1(10):956–64.

10. Chen G, Guo X, Lv F, Xu Y, Gao G. p72 DEAD box RNA helicase is required for optimal function of the zinc-finger antiviral protein. Proc Natl Acad Sci U S A. 2008;105(11):4352–7.

11. Li MM, Lau Z, Cheung P, Aguilar EG, Schneider WM, Bozzacco L, et al. TRIM25 Enhances the Antiviral Action of Zinc-Finger Antiviral Protein (ZAP). PLoS Pathog. 2017;13(1):e1006145.

12. Zheng X, Wang X, Tu F, Wang Q, Fan Z, Gao G. TRIM25 Is Required for the Antiviral Activity of Zinc Finger Antiviral Protein. Journal of virology: J Virol; 2017.

13. Ficarelli M, Wilson H, Pedro Galão R, Mazzon M, Antzin-Anduetza I, Marsh M, et al. KHNYN is essential for the zinc finger antiviral protein (ZAP) to restrict HIV-1 containing clustered CpG dinucleotides. eLife 2019.

14. Buckmaster MV, Goff SP. Riplet Binds the Zinc Finger Antiviral Protein (ZAP) and Augments ZAP-Mediated Restriction of HIV-1. Journal of Virology: American Society for Microbiology; 2022.

15. Gao G, Guo X, Goff SP. Inhibition of retroviral RNA production by ZAP, a CCCH-type zinc finger protein. Science (New York, NY): American Association for the Advancement of Science; 2002. p. 1703–6.

16. Bick MJ, Carroll JW, Gao G, Goff SP, Rice CM, MacDonald MR. Expression of the zinc-finger antiviral protein inhibits alphavirus replication. J Virol. 2003;77(21):11555–62.

17. Liu CH, Zhou L, Chen G, Krug RM. Battle between influenza A virus and a newly identified antiviral activity of the PARP-containing ZAPL protein. Proc Natl Acad Sci U S A. 2015;112(45):14048–53.

18. Mao R, Nie H, Cai D, Zhang J, Liu H, Yan R, et al. Inhibition of hepatitis B virus replication by the host zinc finger antiviral protein. PLoS Pathog. 2013;9(7):e1003494.

19. Muller S, Moller P, Bick MJ, Wurr S, Becker S, Gunther S, et al. Inhibition of filovirus replication by the zinc finger antiviral protein. J Virol. 2007;81(5):2391–400.

20. Tang Q, Wang X, Gao G. The Short Form of the Zinc Finger Antiviral Protein Inhibits Influenza A Virus Protein Expression and Is Antagonized by the Virus-Encoded NS1. J Virol. 2017;91(2).

21. Zhu Y, Chen G, Lv F, Wang X, Ji X, Xu Y, et al. Zinc-finger antiviral protein inhibits HIV-1 infection by selectively targeting multiply spliced viral mRNAs for degradation. Proc Natl Acad Sci U S A. 2011;108(38):15834–9.

22. Miyazato P, Matsuo M, Tan BJY, Tokunaga M, Katsuya H, Islam S, et al. HTLV-1 contains a high CG dinucleotide content and is susceptible to the host antiviral protein ZAP. Retrovirology. 2019;16(1):38.

23. Gonzalez-Perez AC, Stempel M, Wyler E, Urban C, Piras A, Hennig T, et al. The Zinc Finger Antiviral Protein ZAP Restricts Human Cytomegalovirus and Selectively Binds and Destabilizes Viral UL4/UL5 Transcripts. mBio. 2021;12(3).

24. Zhao Y, Song Z, Bai J, Liu X, Nauwynck H, Jiang P. ZAP, a CCCH-Type Zinc Finger Protein, Inhibits Porcine Reproductive and Respiratory Syndrome Virus Replication and Interacts with Viral Nsp9. J Virol. 2019;93(10).

25. Xie L, Lu B, Zheng Z, Miao Y, Liu Y, Zhang Y, et al. The 3C protease of enterovirus A71 counteracts the activity of host zinc-finger antiviral protein (ZAP). J Gen Virol. 2018;99(1):73–85.

26. Trus I, Udenze D, Berube N, Wheler C, Martel MJ, Gerdts V, et al. CpG-Recoding in Zika Virus Genome Causes Host-Age-Dependent Attenuation of Infection With Protection Against Lethal Heterologous Challenge in Mice. Front Immunol. 2019;10:3077.

27. Trus I, Berube N, Jiang P, Rak J, Gerdts V, Karniychuk U. Zika Virus with Increased CpG Dinucleotide Frequencies Shows Oncolytic Activity in Glioblastoma Stem Cells. Viruses: Multidisciplinary Digital Publishing Institute; 2020. p. 579.

28. Udenze D, Trus I, Berube N, Gerdts V, Karniychuk U. The African strain of Zika virus causes more severe in utero infection than Asian strain in a porcine fetal transmission model. Emerging Microbes & Infections 2019; 8:1098–11072019.

29. Darbellay J, Cox B, Lai K, Delgado-Ortega M, Wheler C, Wilson D, et al. Zika Virus Causes Persistent Infection in Porcine Conceptuses and may Impair Health in Offspring. EBioMedicine 25:73–86 2017.

30. Chapagain S, Pal Singh P, Le K, Safronetz D, Wood H, Karniychuk U. Japanese encephalitis virus persists in the human reproductive epithelium and porcine reproductive tissues. PLoS Negl Trop Dis. 2022;16(7):e0010656.

31. Sabir AJ, Singh PP, Trus I, Le NPK, Karniychuk U. Asian Zika virus can acquire generic African-lineage mutations during in utero infection. Emerg Microbes Infect. 2023;12(2):2263592.

32. Darbellay J, Lai K, Babiuk S, Berhane Y, Ambagala A, Wheler C, et al. Neonatal pigs are susceptible to experimental Zika virus infection. Emerg Microbes Infect. 2017;6(2):e6.

33. Trus I, Udenze D, Cox B, Berube N, Nordquist RE, van der Staay FJ, et al. Subclinical in utero Zika virus infection is associated with interferon alpha sequelae and sex-specific molecular brain pathology in asymptomatic porcine offspring. In: Pierson TC, editor. PLoS Pathogens 2019. p. e1008038.

34. Xu MY, Liu SQ, Deng CL, Zhang QY, Zhang B. Detection of Zika virus by SYBR green one-step real-time RT-PCR. Journal of virological methods: J Virol Methods; 2016. p. 93–7.

35. Udenze D, Trus I, Lipsit S, Napper S, Karniychuk U. Offspring affected with in utero Zika virus infection retain molecular footprints in the bone marrow and blood cells. Emerging microbes & infections: Emerg Microbes Infect; 2022. p. 1–43.

36. Bray NL, Pimentel H, Melsted P, Pachter L. Near-optimal probabilistic RNA-seq quantification. Nature Biotechnology. 2016;34(5):525–7.

37. Warren WC, Jasinska AJ, Garcia-Perez R, Svardal H, Tomlinson C, Rocchi M, et al. The genome of the vervet (Chlorocebus aethiops sabaeus). Genome Res. 2015;25(12):1921–33.

38. Law CW, Alhamdoosh M, Su S, Dong X, Tian L, Smyth GK, et al. RNA-seq analysis is easy as 1-2-3 with limma, Glimma and edgeR. F1000Res. 2016;5.

39. The Gene Ontology C. The Gene Ontology Resource: 20 years and still GOing strong. Nucleic Acids Res. 2019;47(D1):D330–D8.

40. Ashburner M, Ball CA, Blake JA, Botstein D, Butler H, Cherry JM, et al. Gene ontology: tool for the unification of biology. The Gene Ontology Consortium. Nat Genet. 2000;25(1):25–9.

41. Raudvere U, Kolberg L, Kuzmin I, Arak T, Adler P, Peterson H, et al. g:Profiler: a web server for functional enrichment analysis and conversions of gene lists (2019 update). Nucleic Acids Res. 2019;47(W1):W191–W8.

42. Todorova T, Bock FJ, Chang P. PARP13 regulates cellular mRNA post-transcriptionally and functions as a pro-apoptotic factor by destabilizing TRAILR4 transcript. Nat Commun. 2014;5:5362.

43. Schwerk J, Soveg FW, Ryan AP, Thomas KR, Hatfield LD, Ozarkar S, et al. RNA-binding protein isoforms ZAP-S and ZAP-L have distinct antiviral and immune resolution functions. Nat Immunol. 2019;20(12):1610–20.

44. Fros JJ, Visser I, Tang B, Yan K, Nakayama E, Visser TM, et al. The dinucleotide composition of the Zika virus genome is shaped by conflicting evolutionary pressures in mammalian hosts and mosquito vectors. PLoS Biol. 2021;19(4):e3001201.

45. Pallett MA, Lu Y, Smith GL. DDX50 Is a Viral Restriction Factor That Enhances IRF3 Activation. Viruses. 2022;14(2).

46. Buechler CR, Bailey AL, Weiler AM, Barry GL, Breitbach ME, Stewart LM, et al. Seroprevalence of Zika Virus in Wild African Green Monkeys and Baboons. mSphere. 2017;2(2).

47. Kerns JA, Emerman M, Malik HS. Positive selection and increased antiviral activity associated with the PARP-containing isoform of human zinc-finger antiviral protein. PLoS Genet. 2008;4(1):e21.

48. Goncalves-Carneiro D, Takata MA, Ong H, Shilton A, Bieniasz PD. Origin and evolution of the zinc finger antiviral protein. PLoS Pathog. 2021;17(4):e1009545.

49. Emeny JM, Morgan MJ. Regulation of the interferon system: evidence that Vero cells have a genetic defect in interferon production. J Gen Virol. 1979;43(1):247–52.

50. Lopusna K, Rezuchova I, Kabat P, Kudelova M. Interferon lambda induces antiviral response to herpes simplex virus 1 infection. Acta Virol. 2014;58(4):325–32.

51. Prescott J, Hall P, Acuna-Retamar M, Ye C, Wathelet MG, Ebihara H, et al. New World hantaviruses activate IFNlambda production in type I IFN-deficient vero E6 cells. PLoS One. 2010;5(6):e11159.

52. Caine EA, Scheaffer SM, Arora N, Zaitsev K, Artyomov MN, Coyne CB, et al. Interferon lambda protects the female reproductive tract against Zika virus infection. Nature Communications 2019 10:1: Nature Publishing Group; 2019. p. 1-12.

53. Lee H, Komano J, Saitoh Y, Yamaoka S, Kozaki T, Misawa T, et al. Zinc-finger antiviral protein mediates retinoic acid inducible gene I-like receptor-independent antiviral response to murine leukemia virus. Proc Natl Acad Sci U S A. 2013;110(30):12379–84.

54. Kozaki T, Takahama M, Misawa T, Matsuura Y, Akira S, Saitoh T. Role of zinc-finger anti-viral protein in host defense against Sindbis virus. Int Immunol. 2015;27(7):357–64.

55. Esser-Nobis K, Aarreberg LD, Roby JA, Fairgrieve MR, Green R, Gale M, Jr. Comparative Analysis of African and Asian Lineage-Derived Zika Virus Strains Reveals Differences in Activation of and Sensitivity to Antiviral Innate Immunity. J Virol. 2019;93(13).

56. Shaw AE, Rihn SJ, Mollentze N, Wickenhagen A, Stewart DG, Orton RJ, et al. The antiviral state has shaped the CpG composition of the vertebrate interferome to avoid self-targeting. PLoS Biol. 2021;19(9):e3001352.

57. Busa VF, Ando Y, Aigner S, Yee BA, Yeo GW, Leung AKL. Transcriptome regulation by PARP13 in basal and antiviral states in human cells. iScience. 2024;27(4):109251.

