## Supplementary figures and images for "Endogenous ZAP is associated with altered Zika virus infection phenotype"

### Figure S1.jpg

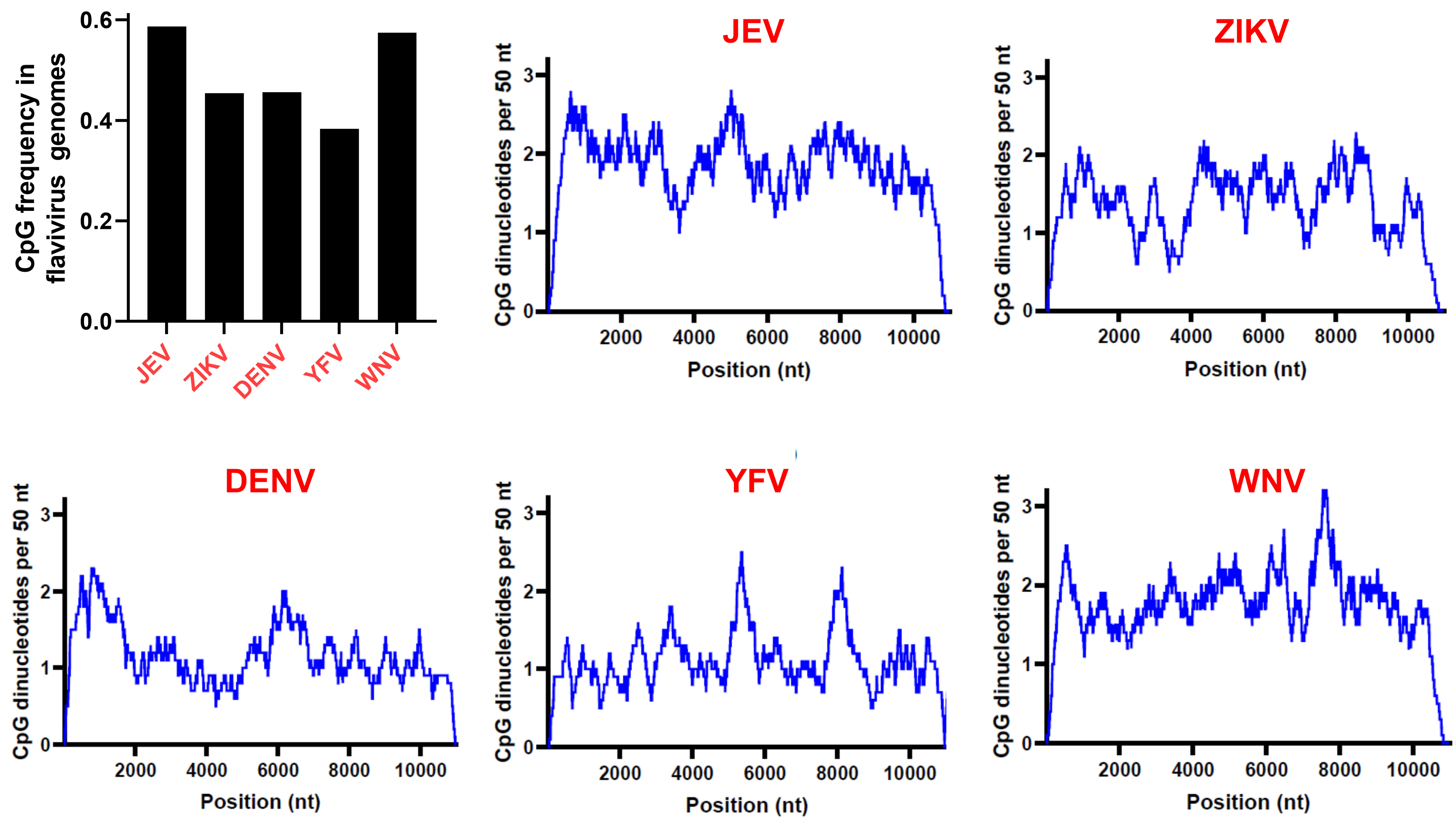

### Figure S2.jpg

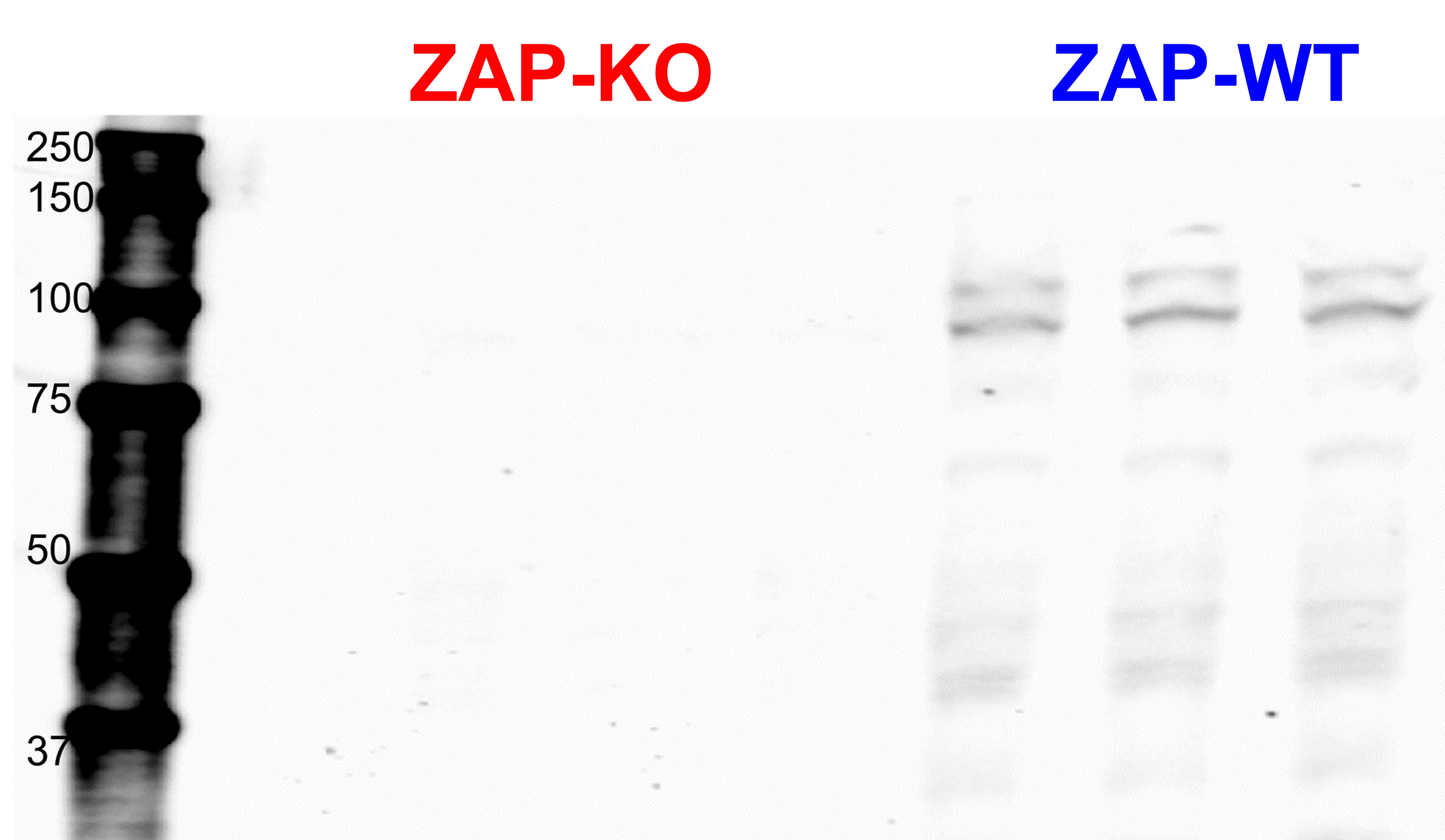

### Figure S3.jpg

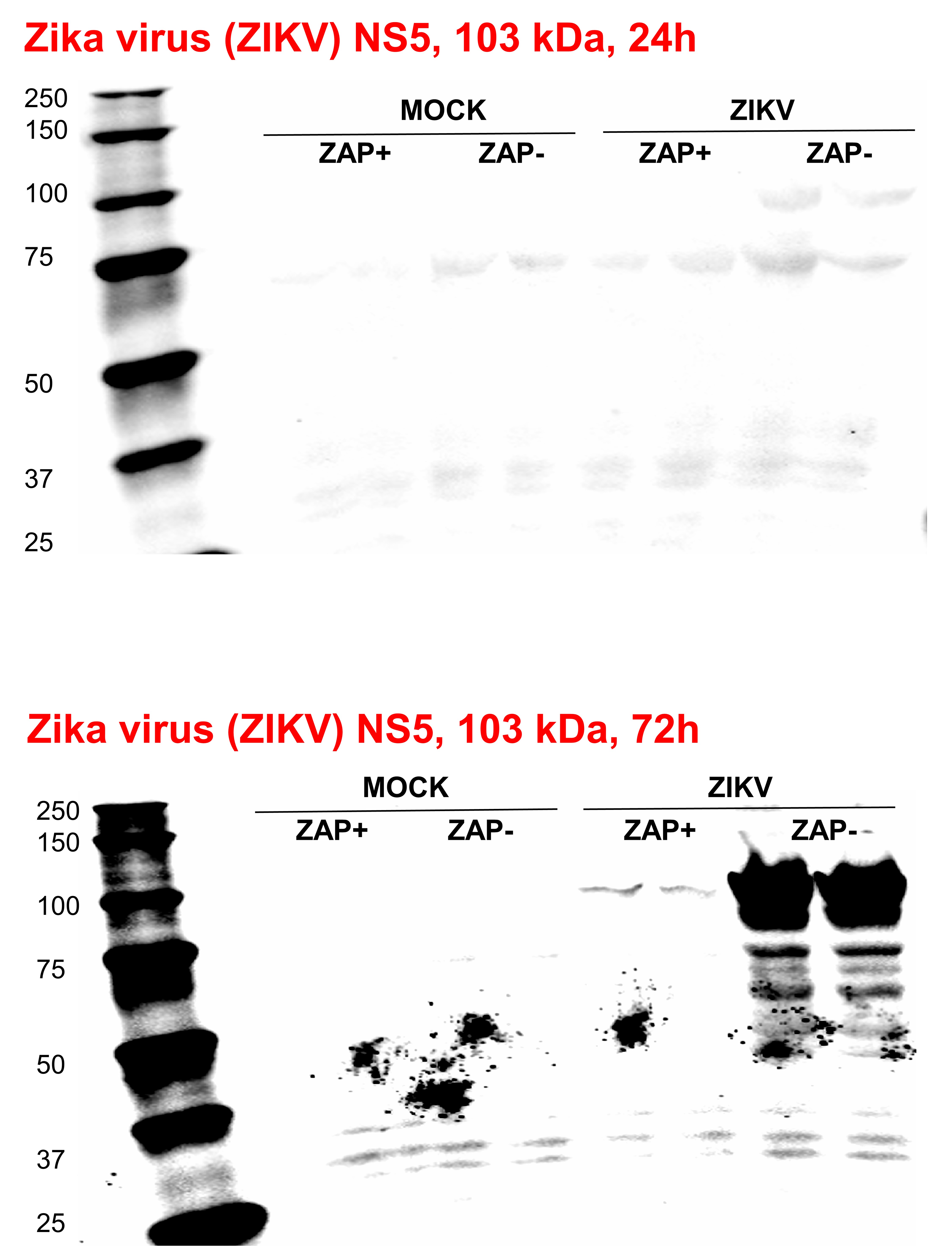
